# Identifying associations between maize leaf transcriptome and bacteriome during different diurnal periods

**DOI:** 10.1101/2025.07.11.664371

**Authors:** Renato Augusto Corrêa dos Santos, Kelly Hidalgo-Martinez, Jorge Mario Muñoz Perez, Daniel Joseph Laspisa, Chenxin Li, Lucas William Mendes, Diego Mauricio Riaño-Pachón, Jason G. Wallace

## Abstract

- Bacterial communities play important roles in the plant phyllosphere. Both microbial communities and their hosts have circadian rhythms and are subject to diurnal environmental changes. However, the interaction between the host and microbiome is still poorly understood. Here, we exploit paired sequencing data of host transcriptome and microbiome derived maize genotypes in field conditions and under two contrasting diurnal periods.
- Gene expression patterns of circadian cycle genes confirmed expected diurnal dynamics. Groups of co-expressed genes that responded to diurnal periods were associated with nucleic acid-binding, heat stress responses, and photosynthesis.
- Microbiome analysis revealed only modest differences in alpha diversity between midday and midnight samples. However, beta diversity indicated a significant shift in community composition. Co-occurrence network analysis identified keystone taxa specific to each time point, suggesting time-dependent ecological roles within the phyllosphere microbiome.
- Cross-correlation analyses between host gene expression and bacterial taxon abundance revealed a greater number of host–microbe associations during the night. Several maize genes involved in circadian regulation significantly correlated with microbial taxa. Our findings provide initial evidence for diurnal-associated relationships between host gene expression and leaf-associated bacteriome, suggesting that maize circadian regulation may play a role in shaping the composition and functional potential of the phyllosphere microbiome.

## Introduction

Endogenous circadian oscillators have been observed and studied in different groups of organisms, including both Bacteria and Plants (Edgar et al. 2012; Greenham and McClung 2015). In plants, the circadian time-keeping mechanism is essential for survival and is important for seasonal processes such as flowering (X. Xu et al. 2022; Harmer 2009). The day-night cycle controls daily and seasonal rhythms (e.g., changes in day length), with diurnal transitions aligning with the internal circadian clock that generates self-sustained rhythms (Hayes et al. 2010). A simplified model of the plant clock, which has largely been studied in *Arabidopsis*, comprises three components: an input pathway sensing light (Zeitgebers for resetting the clock), a core oscillator (transcriptional-translational feedback loops), and output pathways controlling developmental and metabolic processes (physiological adaptations to day-night cycle) (Sanchez and Kay 2016; Harmer 2009; Hayes et al. 2010).

Circadian clocks are associated with important agricultural traits such as increased photosynthesis and growth (Dodd et al. 2005), growth vigor in plants with complex genetics like hybrids and allopolyploids (Ni et al. 2009), leaf senescence (Zhang et al. 2018), seasonal growth, and biotic and abiotic stresses (Bendix, Marshall, and Harmon 2015). Given the importance of grasses as human and animal feedstock and biofuel production, studies to better understand regulatory networks in diurnal gene regulation are needed and can ultimately lead to better strategies for crop improvement. Among grasses, maize (a C4 grass) is an important crop worldwide, and it requires rapid responses to changes in factors such as light, temperature, and humidity over the diurnal cycle (Gao et al. 2023). In both maize and *Arabidopsis*, circadian clock activity is important in the generation of hybrid vigor (Chen 2013; Z. Li et al. 2020). Higher levels of carbon fixation and starch accumulation in maize hybrids are also associated with altered phases of circadian gene expression (Ko et al. 2016).

Integrated analysis of modern sequencing technology datasets has been contributing to improving crops (Yang et al. 2021). In grasses, such studies contribute to our understanding of expression in diurnal cycle genes. For instance, using homologs of *Arabidopsis* circadian genes as a reference, Lai et al. (2020) employed comparative genomics (conserved *cis*-elements) and transcriptomics (RNA-Seq) to examine the conservation and divergence of diurnal gene regulation in C4 grasses sorghum, maize, and foxtail millet (Lai et al. 2020). Another study performed RNA-Seq of maize, from mesophyll and bundle sheath over a 24-hour experiment in constant light and temperature; they recorded increased expression of C4 cycle genes when photosynthesis rates were highest, identified molecular processes associated with temporal changes and employed motif enrichment and co-expression networks to identify regulators in bundle sheath (Borba et al. 2023).

In addition to interest in the host plant genetic or molecular breeding, there is an increasing interest in microbiome engineering of the phyllosphere (i.e., in the upper parts of plants) for crop improvement (Zhan et al. 2022). In the diurnal cycle, the leaf microbiome is subject to environmental variables with light exposure that changes surface moisture, hormone levels, secondary metabolite production, and volatile compound release, which ultimately leads to effects on plant-microbiome interactions (Xu et al. 2022). Therefore, besides understanding host gene regulation in diurnal changes, studies have also been carried out on the composition of the plant microbiome across diurnal cycles. For instance, in a study with a dioecious poplar species, changes in the structure and function of the phyllosphere-associated microbiome were associated with drought tolerance (Lin et al. 2023). The leaf microbiome (endophytic and epiphytic) also harbors a diverse group of nitrogen-fixing bacteria important to plant N supply and growth, and whose activity and composition can be influenced by diurnal cycles (Zhu et al. 2023).

Paired experimental designs in which the host transcriptome and its microbiome are sequenced from the same samples allow to capture associations between the plant transcriptional responses and microbial diversity. For instance, the transcriptional responses of rice seedlings to different soil communities and soil-specific responses were identified (Santos-Medellín et al. 2022). In the C4 cereal sorghum, a large study pairing plant transcriptomics with microbiome (metataxonomics, metagenomics, and metatranscriptomics) data derived from two cultivars in the same experiment provided an enriching resource for studying the association between host metabolism and its microbiome in abiotic stress (drought) (Varoquaux et al. 2019; Xu et al. 2018, 2021). In maize, the most-produced grain on Earth (FAOSTAT, https://www.fao.org/statistics/en), there is still no study analyzing the paired response of the microbiome and the host transcriptome.

In this study, we hypothesized that variations in the plant transcriptome along the diurnal-cycle influence the composition and function of the leaf microbiome, with potential consequences for plant growth and health. To improve our understanding of the potential associations between the host transcriptome and its associated bacteriome in the phyllosphere in contrasting diurnal periods, we used data from two previous studies on maize that independently analyzed the host transcriptome and its bacteriome from a single field experiment (Kremling et al. 2018; Wallace et al. 2018).

## Materials and Methods

### Paired data and preprocessing steps

Paired transcriptome and bacteriome data were obtained from two studies looking at the maize transcriptome (Kremling et al. 2018; Wallace et al. 2018). The original experiment consisted of around 250 maize genotypes, where the upper leaf was sampled ∼2 weeks after flowering time at midday (sample time: 11 AM - 1 PM; ∼30 °C) and midnight (sample time: 11 PM −1 AM; ∼24 °C) on 8 and 26 August 2014. In this study, we used 176 and 228 samples sharing transcriptome and microbiome data, comprising 173 and 222 maize genotypes, for day and night, respectively. RNA from 3 pooled leaf samples per time point was split and processed as described in the respective manuscripts. Illumina sequencing data for the maize transcriptome was obtained from NCBI Bioproject PRJNA383416 (Kremling et al. 2018), which employed 3′ mRNA sequencing (QuantSeq, Lexogen GmBH) with 90-nucleotide single-end reads using Illumina TruSeq primers on an Illumina NextSeq 500 with v2 chemistry. For the present analysis, we filtered RNA-Seq to keep only leaf day and night samples (designated “LMAD” and “LMAN”, respectively, for “Leaf Midpoint At Day/Night”), since those are the only ones for which bacteriome data is also available. The Operational Taxonomic Unit (OTU) table (counts) was obtained from the FigShare repository associated with (Wallace et al. 2018) (https://doi.org/10.6084/m9.figshare.5886769.v2), who sequenced the 16S amplicon to identify bacterial members of the microbial community.

We classified all OTU representative sequences from our previous study (Wallace et al. 2018) based on the Genome Taxonomy Database (GTDB) (Parks et al. 2022) using Qiime2 v. 2024.5 (Bolyen et al. 2019). Briefly, we first trained the qiime2 classifier using rescript (Robeson et al. 2021) and feature-classifier using GTDB v.214 and the fit-classifier-naive-bayes option.

### Functional annotation of the maize genome

Representative CDS and RNA sequences associated with primary transcripts of maize NAM genome version 5 (Hufford et al. 2021) were obtained from Phytozome (version Zmays_Zm_B73_REFERENCE_NAM_5_0_55). Functional annotation was carried out using InterProScan 5 (Jones et al. 2014) to obtain InterPro domains and GO terms. Transcription Factors (TFs) were identified using HMMER 3 (Mistry et al. 2013) and pre-defined rules for the identification of Transcription-Associated Proteins (TAPs), including TFs (Pérez-Rodríguez et al. 2010).

### Transcriptome and co-expression network analysis

Leaf day & night transcriptome data were pre-processed to quantify gene expression. Briefly, we checked the quality of raw Illumina reads using FastQC (Andrews, 2010). Cleaning steps were carried out by TrimGalore v.0.6.10 (Krueger et al. 2023). Salmon v.1.9 (Patro et al. 2017) was used to quantify transcript expression (index and quant options). Final matrices were generated with Salmon option quantmerge, with estimated read counts and normalized expression data (transcript per million, or TPM).

Since we intend to compare gene expression in different diurnal periods (midday and midnight), we inspected the expression patterns using a Principal Component Analysis (PCA) using the R prcomp() function from the stats library (R Core Team 2024). Additionally, we inspected a few of the maize circadian genes (**Table 1**), previously described (Lai et al. 2020), by generating violin plots of their TPM expression values.

**Table 1.** Genes of the circadian cycle with expected expression peaks close to the middle of day or night (based on Lai et al 2020; EF3L2 was extracted from Zhao et al 2023)

| Original gene identifier | Maize v5 | Expected expression profile |
| --- | --- | --- |
| GRMZM2G045275 - ELF3-LIKE 1 (EF3L1) | Zm00001eb159890_T001 | Higher expression - night |
| GRMZM2G095727 - PRR73-LIKE (P73L) | Zm00001eb397240_T001 | Higher expression - day |
| GRMZM2G421256 - RVE7-LIKE 3 (RE7L3) | Zm00001eb072870_T003 | Higher expression - night |
| GRMZM2G181030 - RVE7-LIKE 4 (RE7L4) | Zm00001eb428070_T001 | Higher expression - night |
| ELF3-LIKE 2 (EF3L2) | Zm00001eb297250_T001 | Higher expression - night |

Before reconstruction of the co-expression network with both day and night samples (n = 404), we filtered out genes with low expression by keeping only genes with at least 3 RPKM in over 80% of samples. We also filtered by the coefficient of variation (standard deviation/mean) by keeping only genes over the first quartile. These matrices are also used in the cross-correlation analyses involving the bacteriome.

A co-expression network was reconstructed for the day and the night samples using corALS (Becker et al. 2023) and computing Pearson correlation coefficients (PCC). Network edges were filtered by keeping only gene pairs with PCC > 0.7 (only positive correlations considered), followed by clustering with the Leiden algorithm as implemented in the igraph R package (Csárdi et al. 2025). Optimal resolution for module detection followed the strategy described in the “Simple Tidy GeneCoEx” R workflow (Li, Deans, and Buell 2023), which maximizes the number of modules and genes, considering a minimum module size of 5 genes. Eigengenes of expression values in co-expression modules responding to midday - midnight periods were computed for heatmap visualization using the pheatmap R package (Kolde, 2025). We used the find_enrichment.py script from GOATOOLS (Klopfenstein et al. 2018) to identify enriched GO terms (MF, BP, and CC) in each module.

### Bacteriome and co-occurrence network analysis

The original OTU counts matrix generated by (Wallace et al. 2018) was used in downstream analyses and was generated from amplicon sequencing (16S) from the total RNA sample, in order to capture metabolically active bacteria. Only bacteriome samples with a corresponding transcriptome from the same sample (plot and time point) were kept. OTUs were kept if their relative abundance was over 0.001% in at least 50% of total samples with transcriptome data (n = 404) and with the coefficient of variation over the first quartile.

We performed the weighted unifrac analysis in QIIME 2 v2023.7 (Bolyen et al. 2019). Briefly, using the filtered subset of OTUs, we aligned the representative 16S rRNA sequences with MAFFT (Katoh and Toh 2008) and inferred a phylogeny with IQ-TREE (Minh et al. 2020). The tree was rooted at the midpoint. We used the ‘qiime diversity core-metrics-phylogenetic’ function to compute beta diversity metrics using a sampling depth of 1,000 reads (--p-sampling-depth option). Tests of differences between groups based on alpha diversity were carried out using the ‘qiime diversity alpha-group-significance’ for evenness, Faith’s PD, and Shannon metrics. PERMANOVA for differences between diurnal periods was computed with ‘qiime diversity beta-group-significance’ using the Weighted Unifrac distance matrix with 999 permutations. Differential Abundance Analysis (DAA) was computed using the “Analysis of Compositions of Microbiomes with Bias Correction” - ANCOMBC method (Lin and Peddada 2020).

Networks were reconstructed with the Spiec-Easi method (Kurtz et al. 2015) using options method=’mb’ (Meinshausen-Bühlman), lambda.min.ratio=1e-2, nlambda 20, pulsar.params (rep.num 50). We clustered the networks using the Louvain algorithm (cluster_louvain) implemented on the igraph R package (Csárdi et al. 2025), with the default resolution (1). To make the analysis reproducible, we used the set.seed() R function along with the “L’Ecuyer-CMRG” random number generator in network generation and clustering steps.

To identify potential keystone species in the network, within-module (Zi) and among-module connectivity (Pi) metrics were computed using the within_module_deg_z_score and part_coeff function from the brainGraph R library (Watson 2015). Nodes were defined as peripherals/specialists (Zi < 2.5; Pi < 0.62), module hubs (Zi ≥ 2.5; Pi < 0.62), network hubs (Zi ≥ 2.5; Pi ≥ 0.62), and connectors (Zi < 2.5; Pi ≥ 0.62) (Olesen et al. 2007). An enrichment analysis was carried out to identify enriched genera in groups of potential keystone taxa, using GTDB (version 214) assignments. We employed the universal enrichment analyzer as implemented in the enricher function from clusterProfiler (Yu et al. 2012) to identify enriched species in each group of keystone taxa, using OTUs from the filtered dataset as background genes and GTDB genera in term2gene mapping.

### Cross-correlation between transcriptome and bacteriome

Bacteriome data generated and analyzed by (Wallace et al. 2018) comprises a subset of samples analyzed by (Kremling et al. 2018), opening the possibility of paired transcriptome-bacteriome analysis for samples present in both studies. To compute cross-correlations between the host transcriptome and bacteriome, we used estimated expression raw counts from Salmon and the OTU counts abundance after filtering steps, as described in their respective sections above. Cross-correlations for the day and the night datasets were computed with SparXCC (Sparse Cross-Correlation between Compositional data) (Jensen et al. 2024), using pseudo_count = 1, var_min = 1e-05, Find_m option to enable dynamic search for permutation threshold (m), and 100 bootstrap replicates to calculate m.

To explore the potential functions associated with OTUs that were significantly cross-correlated with the host transcriptome, we performed a putative functional prediction with PICRUST2 v.2.4.1 (Douglas et al. 2020). Functional enrichment analysis of predicted KEGG Orthologs (KOs) was conducted using the enricher function from clusterProfiler (Yu et al. 2012).

## Results

### Co-expression network analysis reveals different gene modules responding to diurnal periods in maize leaves

In this work, we investigated the association between host bacteriome and transcriptome in maize leaves using public sequencing data obtained from a large field experiment carried out previously (Wallace et al. 2018; Kremling et al. 2018). Given the availability of day and night samples, the consequent expected variation in maize gene expression across the diurnal cycle (Lai et al. 2020; Hayes et al. 2010) and in metabolically active bacteria (Gunnigle et al. 2017; Newman et al. 2022) due to changes in light and temperature, we re-analyzed these two omics datasets independently as well as their associations using network approaches with particular focus on contrasts between day and night.

We re-analyzed RNA-Seq data from Kremling et al. (2018); however, since our study aims to investigate host transcriptome and bacteriome, we only used the RNA-Seq datasets derived from samples with paired amplicon data (metataxonomics) in Wallace et al. (2018). This subset of samples comprises 176 and 228 samples for day and night diurnal periods, respectively. Except for the B73 maize line (which was used as a replicated check line), samples correspond to different field plots, and each represents a different maize genotype.

We used stringent filtering steps based on minimum expression across samples (at least 3 RPKM in > 80% of samples) and keeping genes with a coefficient of variation above the lowest quartile, which reduced from approximately 40k to 7,201 genes. We first checked the overall expression patterns to ensure that a comparison between diurnal periods is feasible. Principal Component Analysis (PCA) showed a clear separation between day and night samples (**Figure 1A**); in contrast, we did not observe such separation when comparing expression across maize subpopulations (**Sup Fig S1**). The expression of selected maize circadian cycle genes expected to peak around midday or midnight (Lai et al. 2020) was also consistent, except for one gene (ELF3-LIKE 1 - EF3L1, Zm00001eb159890_T001) (**Figure 1B**). However, a homolog (ELF3-LIKE2) showed the expected pattern and may be sufficient for its role.

**Figure 1.**
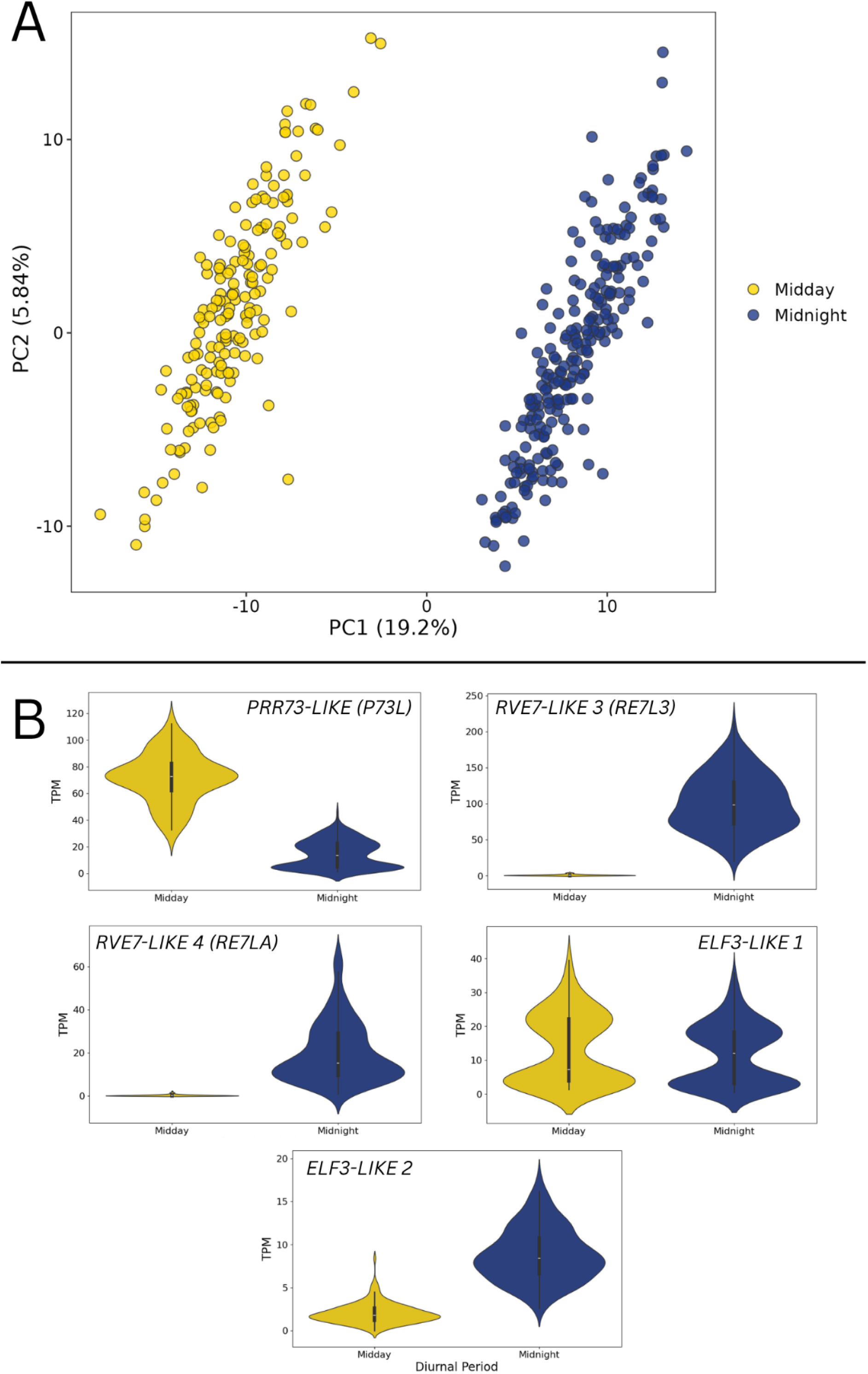
Exploratory analysis of the original dataset in the context of diurnal period comparison. **(A)** Principal Component Analysis (PCA) based on gene expression highlighting diurnal periods. **(B)** Gene expression for selected circadian genes with expression peak around midday (PRR73-LIKE, P73L) or midnight (RVE7-LIKE 3, RE7L3; and RVE7-LIKE 4, RE7L4) (Lai et al. 2020). Except for ELF3-LIKE 1 (Zm00001eb159890_T001), all others in Lai et al (2020) showed the expected pattern of day/night expression in our analyses. ELF3-LIKE 2 (Zm00001eb297250_T001) is a homolog of ELF3-LIKE 1, described in Zhao et al (2023), which has the expected expression pattern.

To identify genes involved in response to diurnal periods and their associated functions, we used a network approach. We first reconstructed a co-expression network including the RNA-Seq data for the subset of 7,201 genes remaining after filtering steps described previously. For this step, we computed Pearson correlation coefficients (PCC) for all pairs of genes using corALS (Becker et al. 2023). We filtered out low correlations, keeping only PCC ≥ 0.7. Among all co-expression modules (33 total), we identified eighteen modules showing significant differences between day and night (**Sup Fig S2; Sup Table S1**).

To investigate functions associated with modules, we used the GO annotation of maize genes and GOATOOLS to find enriched terms. We found enriched terms in three modules (**Table 2; Figure 2**). Interestingly, module #1 (205 genes) is enriched in nucleic acid binding, and has several transcription factors (AP2-EREBP, C2C2-CO-like, C2C2-GATA, C2H2, CCAAT, FAR1, GRAS, HB, MYB-related, and WRKY), and has a higher expression during midnight. Module #5 (67 genes) is likely associated with heat stress response, and it has a higher expression during midday. Module #9 (45 genes), also with higher expression during midday, showed significant terms associated with photosynthesis and energy generation, with no associated TFs. These results highlight groups of genes that are responding to changes in diurnal periods (both circadian and diel responses), some of which may be associated with the changing bacteriome.

**Figure 2.**
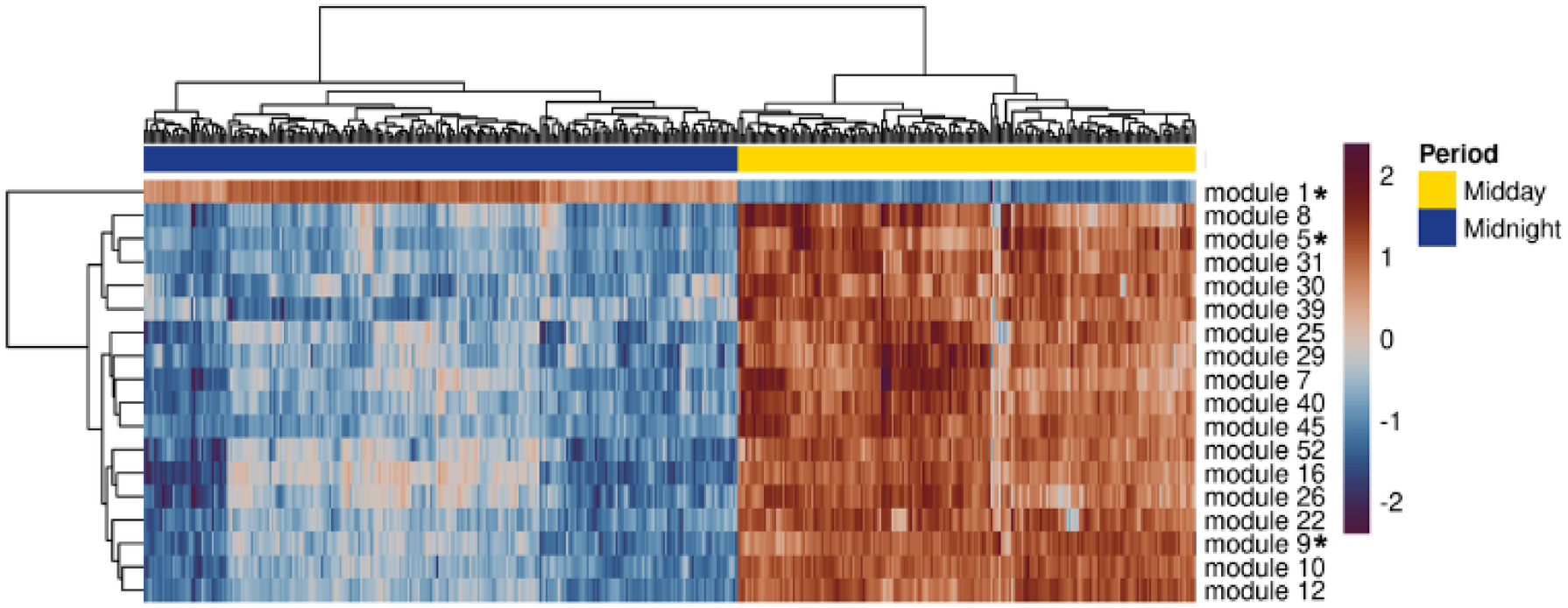
Relative expression value of eigengenes (z-score) of the eighteen co-expression modules with contrasting expression between midday and midnight. Three marked modules (*) are enriched for Nucleic Acid Binding (1), Heat Stress Response (5) and Photosynthesis (9).

**Table 2.** GO enrichment analysis of co-expression modules with expression profiles of contrasting diurnal periods.

| Module | Gene Ontology term | Significance (Bonferroni) | Diurnal period higher expression |
| --- | --- | --- | --- |
| 1 | GO:0003676 - nucleic acid binding | 0.022 | Night |
| 5 | GO:0044183 - protein folding chaperone | < 1E-30 | Day |
|  | GO:0051082 - unfolded protein binding | < 1E-30 |  |
|  | GO:0140662 - ATP-dependent protein folding chaperone | < 1E-30 |  |
|  | GO:0031072 - heat shock protein binding | 0.01 |  |
|  | GO:0005515 - protein folding | < 1E-30 |  |
| 9 | GO:0009765 - photosynthesis, light harvesting | 9.23E-25 | Day |
|  | GO:0006091 - generation of precursor metabolites and energy | 1.55E-12 |  |
|  | GO:0015979 - photosynthesis | 0.01 |  |
|  | GO:0008152 - metabolic process | 0.04 |  |
|  | GO:0016020 - membrane | 7.85E-08 |  |
|  | GO:0110165 - cellular anatomical structure | < 1E-30 |  |
|  | GO:0005575 - cellular_component | 0.01 |  |
|  | GO:0009521 - photosystem | 0.04 |  |

### Potential keystone bacterial species differ among diurnal periods

To understand potential functions in the maize leaf bacteriome associated with the host transcriptome, we used the Operational Taxonomy Unit (OTU) matrix available from Wallace et al. (2018), which comprises approximately 9K OTUs. We updated the OTU classification by training a qiime2 classifier to assign identifiers to OTUs based on the GTDB database (Parks et al. 2022) using their 16S rRNA representative sequences (**Sup Table S2**). Similarly to host transcriptome analyses, we applied filters to keep only those OTUs with significant abundance and variability. We removed OTUs that showed a relative abundance below 0.001 in at least half of the samples and that had a coefficient of variation below the first quartile. This left 276 OTUs, corresponding to 74.67% of total OTU read counts (samples with a paired RNA-Seq).

This subset of OTUs mostly comprises members of the orders Sphingomonadales, Rhizobiales, and Actinomycetales, even though at the genus level most OTUs have no taxonomic assignment (**Sup Fig S3**). Similarly to the transcriptome data, a Weighted Unifrac analysis showed separation (albeit less strong) of day and night samples (**Figure 3**), and PERMANOVA tests resulted in significant differences (p-value = 0.001) between diurnal periods. Tests of differences between diurnal periods based on alpha diversity were not significant (**Sup Fig S4**).

**Figure 3.**
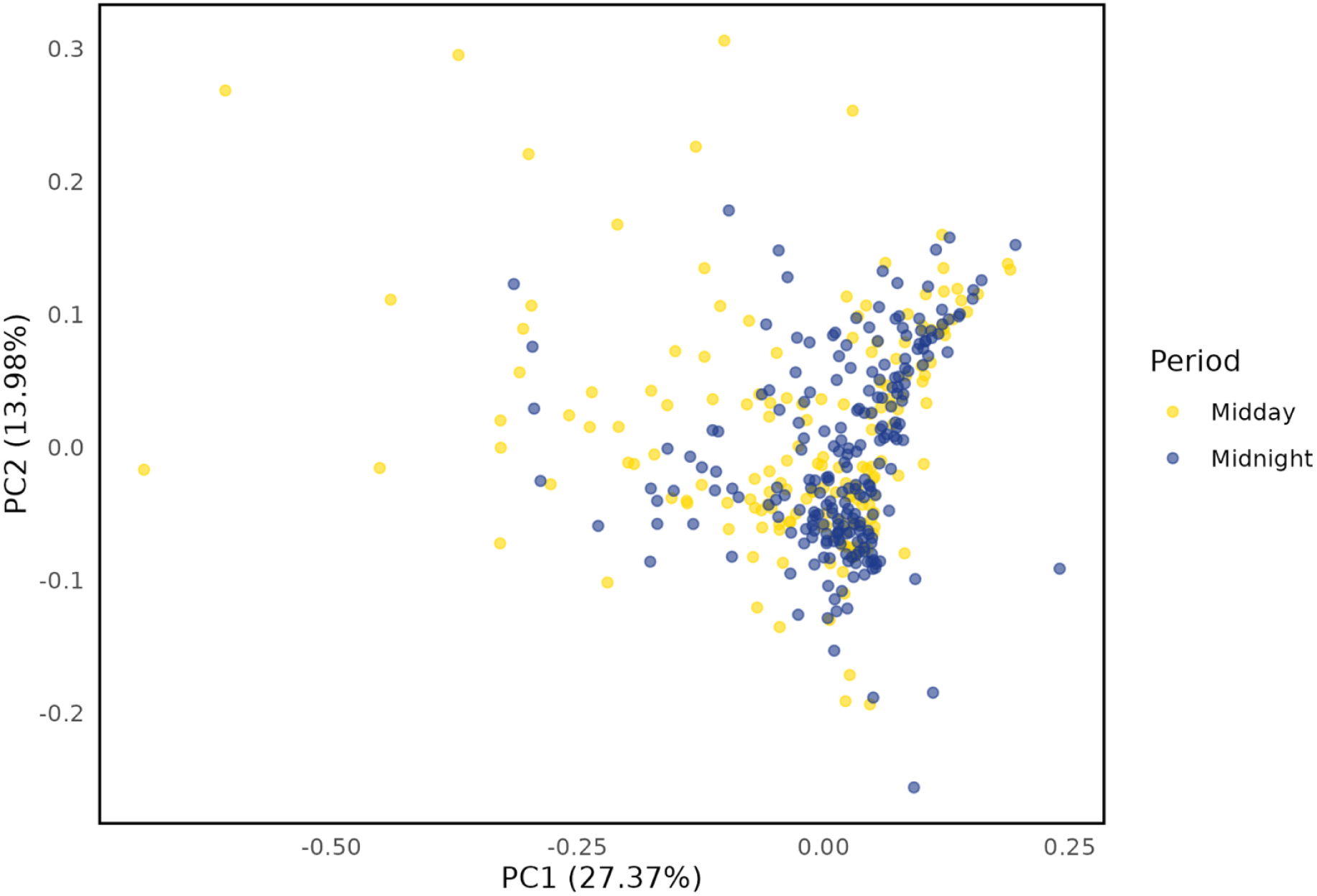
Beta-diversity analyses of the bacteriome data (Weighted Unifrac), showing the first two components. PERMANOVA analysis of samples indicates that day and night samples are significantly different from each other (p=0.001).

We used the ANCOMBC method to identify differentially abundant OTUs, detecting 36 more abundant at midday and 51 at midnight (**Figure 4; Sup Table S3**). Of the ones more abundant at night, eight of the top 10 OTUs with the highest differential abundance are classified as *Sphingomonas* and/or Sphingomonadaceae, plus one *Hymenobacter* sp. and an unclassified member of the family Microbacteriaceae. OTUs with more abundance during the day are more diverse, with the top 10 including members of families Enterobacteriaceae, Burkholderiaceae, and Sphingomonadaceae, and species of *Methylobacterium*, *Pedobacter*, *Chryseobacterium*, *Agrobacterium,* and *Brevundimonas*. Interestingly, 82% of the 51 OTUs more abundant at night (=42 total) correspond to either *Sphingomonas* spp. or an unassigned member of Sphingomonadaceae, whereas only 36% (13) of day-biased OTUs are in this family.

**Figure 4.**
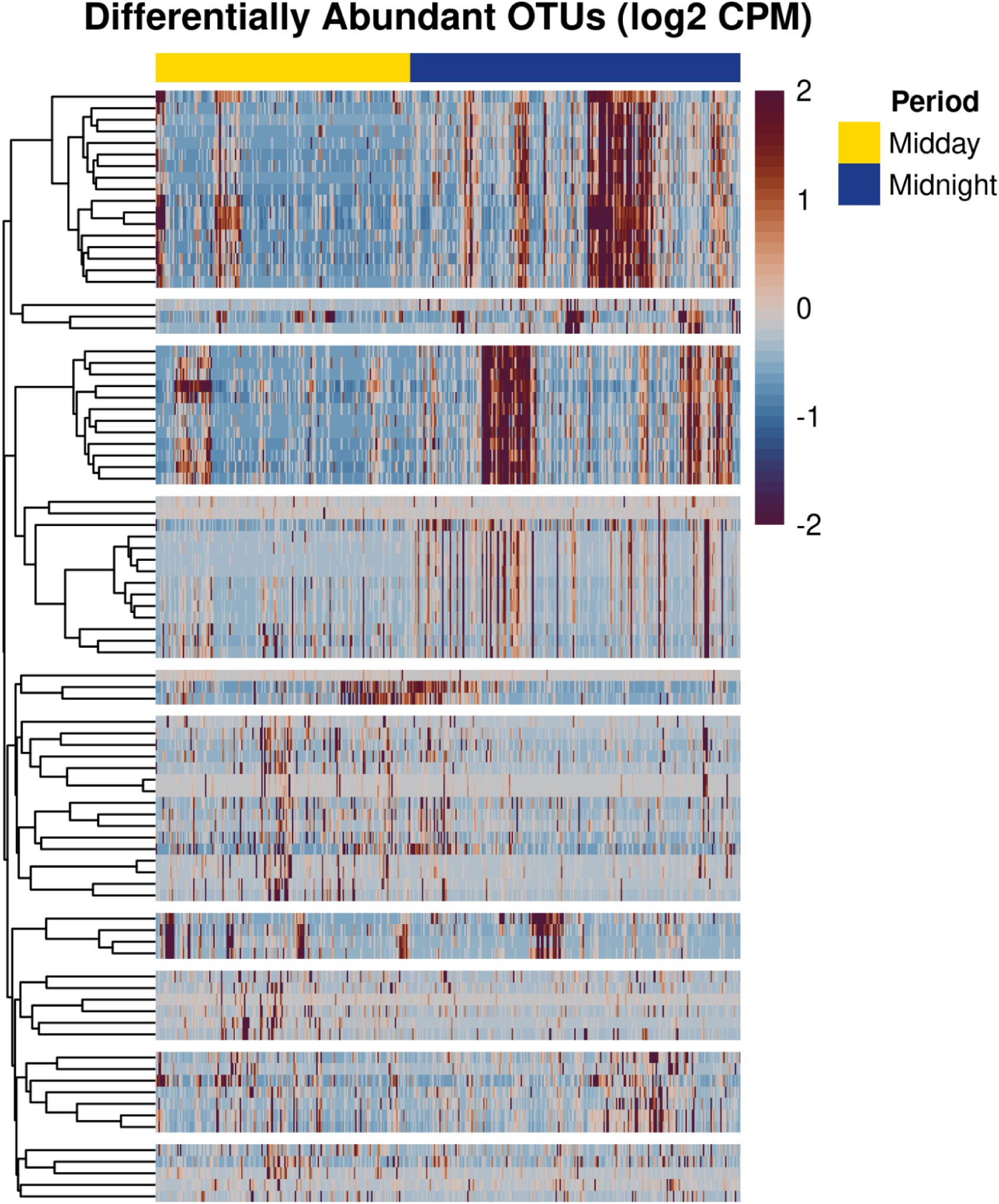
Heatmap of all differentially abundant OTUs between diurnal periods (87 total, y axis), showing relative abundance (z-score) and grouped by diurnal period (x axis). OTUs were clustered with the R pheatmap() function with default settings (complete linkage and euclidean distances).

Since functional inference based on 16S rRNA alone is difficult, we employed a network approach to identify the potential ecological roles of nodes (OTUs) in the network. We were particularly interested in the functional cartography of the network, by identifying module and network hubs as well as connectors, collectively called keystone species (Guimerà and Nunes Amaral 2005). For each diurnal period, we reconstructed co-occurrence networks using the SpiecEasi algorithm (Kurtz et al. 2015), which uses a conditional dependence graphical approach and tends to represent associations more accurately than correlations. The day and night networks have similar modularity, with seven and nine modules, respectively (**Figure 5A-B; Table 3; Sup Table S4**). Assuming network nodes with similar topological properties have similar functional roles (Guimerà and Nunes Amaral 2005), we employed a clustering method followed by computation of the within- and among-module connectivity metrics to identify OTUs representing peripherals (which have a low number of connections inside and among modules), potential module hubs (which have a high number of connections inside network modules), connectors (which have a high number of connections among modules), and network hubs (which have a high number of connections both within and among modules).

**Figure 5.**
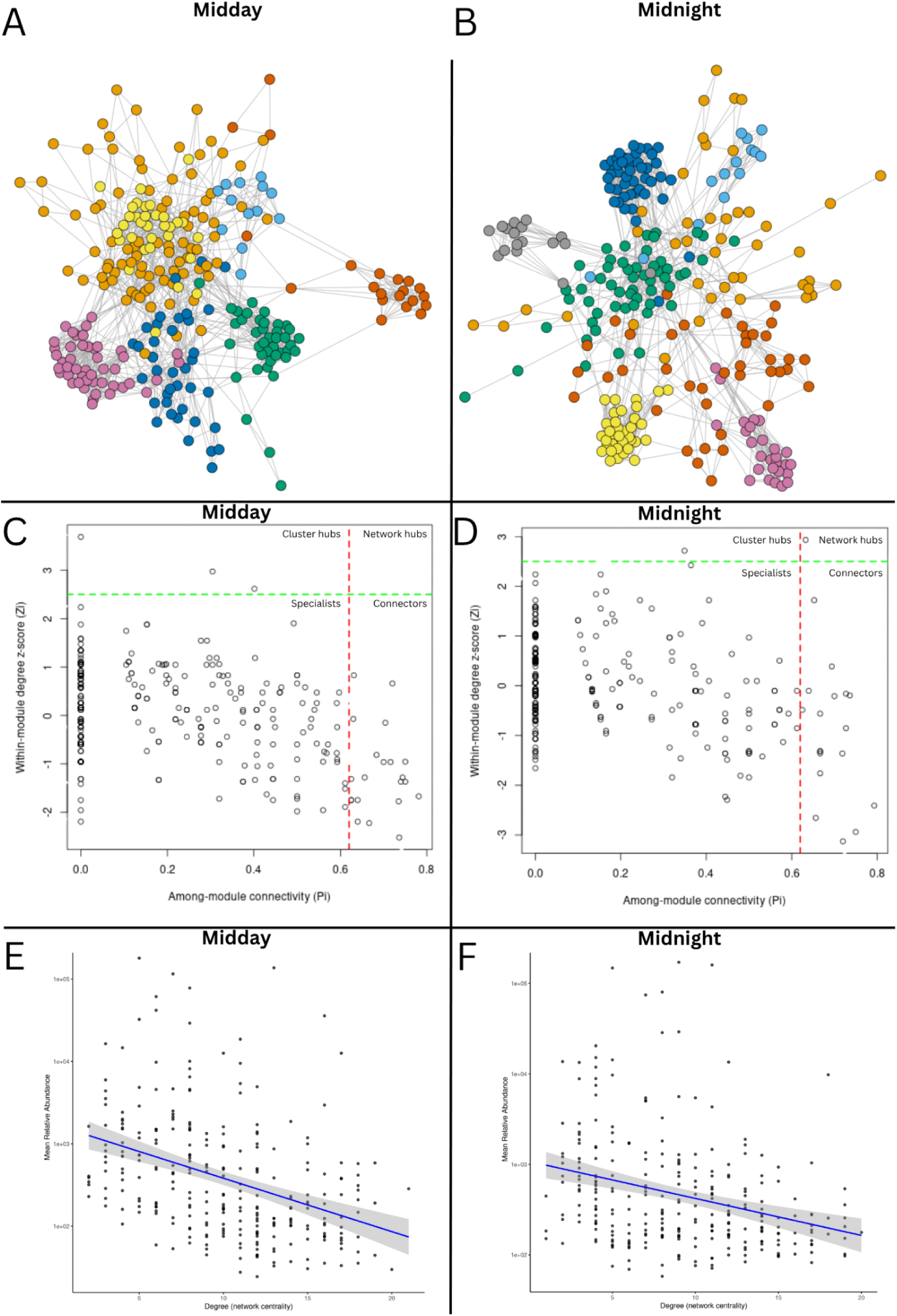
Co-occurrence networks for the day (left) and the night (right) samples. **(A-B)** Co-occurrence networks, with colors indicating different Louvain modules. **(C-D)** Potential keystone species identified using the Pi (x axis) and Zi (y axis) metrics. Connectors have high Pi scores, module hubs have high Zi scores, and network hubs are high for both metrics; dashed lines indicate the cutoffs for each metric. **(E-F)** Relationship between node degree and mean CPM values in the subsets (276 OTUs).

**Table 3.** Metrics of co-occurrence networks in both diurnal periods (day and night)

| **Network Metric** | **Day** | **Night** |
| --- | --- | --- |
| ***Number of modules (Louvain )*** | 7 | 9 |
| ***Number of edges*** | 1,388 | 1,255 |
| ***Number of positive edges*** | 1,234 | 154 |
| ***Number of negative edges*** | 1,179 | 76 |
| ***Average degree*** | 10.06 | 9.09 |
| ***Clustering coefficient*** | 0.31 | 0.37 |
| ***Centrality closeness*** | 0.17 | 0.17 |
| ***Centrality betweenness*** | 0.06 | 0.08 |

We noticed that both day and night networks have most nodes assigned to potential specialists (nodes having few connections to members of different modules) (**Figure 5C-D; Sup Table S4**). By analyzing potential keystone taxa corresponding to nodes with an increased number of connections with other nodes in day and night networks, we noticed that just one network hub was identified at midnight, comprising a member of Enterobacteriaceae, and no network hub at midday. Both networks have just a few module hubs, including a shared species of *Deinococcus*, and a larger number of connectors, the highest group of keystone species in both networks. Interestingly, taxa with higher network degree tend to have lower abundance profiles compared to those with lower degree, suggesting that rare taxa may have stronger effect on network structure (**Figure 5E-F**). Most day and night connectors are *Sphingomonas* and *Methylobacterium* species. We also carried out an enrichment analysis to identify over-represented genera in groups of keystone species, but no significant enrichment was detected in any group.

### Associations between gene expression and potential keystone species differ between diurnal periods

To identify potential associations between members of the microbial communities and the host-expressed genes, we employed SparXCC, which computes cross-correlations between pairs of OTUs and genes. For this analysis, we used the filtered subset of maize genes (7,001 genes) and OTUs (276 OTUs) used for co-expression and co-occurrence networks, respectively; however, differently from the transcriptome and microbiome analyses, in which we used a different number of samples for midday (n = 176) and midnight (n = 228), in cross-correlations we retained only samples derived from field plots having both transcriptome and bacteriome for both midday and midnight periods. In total, 174 plots were used for this analysis.

Using a dynamic selection for the permutation threshold, which defines a minimum value for correlations to be considered “significant”, SparXCC identified 650 correlations in the day samples (290 genes; 76 OTUs) and 1,700 correlations in the night samples (732 genes; 83 OTUs), including both positive and negative correlations (**Sup Fig S5; Sup Table S5**). Two of the circadian clock or clock-associated maize genes showed cross-correlations with OTUs (Zm00001eb122110_T003, GIGANTEA 2-GI2; Zm00001eb397240_T001, PRR73-LIKE-P73L), mostly showing negative correlations. Among genes in cross-correlations with the bacteriome, we identified members of transcription factor families in midday (13/290) and midnight (54/732) (**Sup Table S6**).

To investigate potentially important microbes and associated gene functions in the day and night networks, we selected edges with the absolute value of cross-correlations > 0.5. Interestingly, the OTU associated with *Deinococcus,* also considered a module hub in co-occurrence network analyses, showed strong positive and negative correlations with genes in both networks (including additional significant cross-correlations > 0.5). However, the exact groups of genes associated with *Deinococcus* change across both networks.

To further investigate functions associated with correlated genes in the network, we tested GO and KO enrichment for genes and OTUs, respectively, in the whole day and night networks, using GOATOOLS. Interestingly, a GO term associated with photosynthesis (“photosynthesis, light harvesting” - GO:0009765) was over-represented in host gene sets for both diurnal periods (day adj. p-value = 0.00004; night adj. p-value = 0.01324). For the OTUs, both diurnal periods showed enriched KOs of ribosomal proteins (e.g., rplW, rpsJ, rpsD, rpsF, and rpsG), conjugal transfer pilus assembly proteins (e.g., traG, traB, traF, traU,traW, traK, and traE), and transporters (fadL, tetA, TC.ZIP, TC.HAE1, zntA, TC.BAT1, and copA). Day and night-specific enriched terms were also identified.

One of the modules identified in the clustered co-expression network also showed significantly enriched terms associated with photosynthesis and energy production (module #9). Interestingly, 14 and 18 genes from that module were also identified in the cross-correlations of host genes and bacterial OTUs in day and night samples, respectively, including 7 shared genes.

## Discussion

No previous study has investigated the associations between the host transcriptome and bacteriome in the maize phyllosphere, or how these relationships vary across different diurnal periods. In this work, we analyzed publicly available datasets from a large-scale field experiment involving diverse maize genotypes to explore host transcriptome–bacteriome associations in maize leaves at two contrasting time points: midday and midnight.

We started by examining transcriptomic changes in maize leaves between midday and midnight. Early studies using microarrays revealed thousands of genes responsive to the diurnal cycle in maize leaves, in contrast to non-photosynthetic tissues, where relatively few genes, including core oscillator genes, exhibited diurnal regulation (Hayes et al. 2010). More recently, a comparative analysis across grasses found that approximately 30% of maize leaf transcripts display diurnal rhythmicity (Lai et al. 2020). In addition, Borba et al. (2023), analyzing gene expression across the day in mesophyll and bundle sheath cells, identified cell type and time of day as the primary drivers of variance in their dataset. These previous results are consistent with our observations, as we also detected a strong separation between day and night transcriptomic profiles.

Focusing on circadian clock genes expected to show contrasting expression between midday and midnight, we confirmed that three genes displayed the anticipated diurnal patterns. Interestingly, a member of the Evening Complex (composed of ELF3, ELF4 and LUX, a MYB-domain transcription factor) (Zhao et al. 2023; Lai et al. 2020), ELF3-like (Zm00001eb159890) did not show the expected higher expression at night, although its homolog *ZmELF3.2* (Zm00001d039156) did, suggesting that this homolog may be functionally compensating in maize.

Co-expression network analysis revealed multiple gene modules associated with diurnal variation, including modules enriched for ‘nucleic acid binding’ (higher at night), ‘photosynthesis’ (higher during the day), and ‘heat response’ (also higher during the day). Nutrient demand changes with physiological alterations as a consequence of diel alterations, including photosynthesis and transpiration rates (Haydon, Román, and Arshad 2015). The circadian clock is known to be associated with biotic and abiotic stress responses, including drought and high or low temperature [reviewed in (Sanchez and Kay 2016)]; therefore, given temperature differences observed in our experiments (delta of approx. 6 °C between midday and midnight), it was also expected to observe groups of genes associated with heat response.

The phyllosphere environment is complex and variable, with rapid changes in aspects such as water content, nutrients, temperature, and light exposure. Microbial communities inhabiting this niche can originate from diverse sources—including soil, air, and neighboring plants, and their structure is influenced by multiple factors such as host genotype, seasonality, and environmental conditions (Vorholt et al. 2012). Despite this complexity, the phyllosphere of seed plants is usually dominated by a few genera, such as *Sphingomonas*, *Methylobacterium,* and *Pseudomonas* (Sohrabi et al. 2023). In our dataset, *Sphingomonas* and *Methylobacterium* were consistently among the most abundant taxa. Interestingly, *Methylobacterium* species have been highlighted as an important Plant Growth Promoting Bacterium (PGPB) by producing indole acetic acid (Palberg et al. 2022). Also, both are genera with members of diazotrophic bacteria (= nitrogen fixing) that are typically identified in the phyllosphere (Zhu et al. 2023).

Previous studies have provided evidence for associations between the plant diurnal cycle and the structure of associated microbial communities. For example, Zhao et al. (2021) demonstrated that microbial abundances in the rice rhizosphere shift between day and night under both light–dark cycles and constant darkness, with reduced diversity observed under daytime conditions. Similarly, Newman et al. (2022) showed that disruptions in the host circadian rhythm of *Arabidopsis thaliana* altered the temporal dynamics of rhizosphere bacterial communities. Additional studies in the rhizospheres of *A. thaliana* and *Brachypodium distachyon* also reported significant day–night differences in microbial composition (Staley et al., 2017). Despite these insights from root-associated microbiomes, similar investigations in the phyllosphere remain scarce. In our study, alpha diversity did not differ significantly between diurnal time points. However, PERMANOVA based on Weighted UniFrac distances revealed significant differences in community structure, suggesting that the phyllosphere microbiota is phylogenetically distinct between midday and midnight.

Co-occurrence networks are widely used to infer potential associations among microorganisms, including putative ecological interactions. Several computational methods have been developed to deal with properties such as compositionally and sparsity that are associated with microbiome amplicon sequencing data (Matchado et al. 2021). Assuming that nodes with similar topological properties tend to have similar roles (Banerjee, Schlaeppi, and van der Heijden 2018; Guimerà and Nunes Amaral 2005), we employed network metrics such as degree and centrality to identify specialists and potential keystone species in maize day and night co-occurrence networks.

In both midday and midnight co-occurrence networks, most nodes were classified as peripherals, consistent with patterns observed in previous studies (Pan et al. 2021; Hidalgo-Martinez et al. 2024). Among the taxa identified as potential keystones, only a few were predicted to function as module hubs. Notably, a *Deinococcus* species was identified as a shared module hub in both networks. *Deinococcus* has also been reported as a keystone taxon in various microbiomes in previous studies (Zhu et al. 2021).

In our analysis, one OTU assigned to the genus *Pseudomonas* was a module hub in the day network. *Pseudomonas* species have previously been identified as keystone taxa in soil aggregate microbiomes (Xiong et al. 2021). In the phyllosphere context, *Pseudomonas putida*, in combination with *Stenotrophomonas maltophilia*, has been shown to promote plant growth in maize leaves, leading to physiological changes such as increased chlorophyll content, likely contributing to enhanced photosynthesis (Abadi et al. 2020).

Several OTUs assigned to *Sphingomonas* and *Methylobacterium* were identified as connectors in both midday and midnight networks. Our analysis of this phyllosphere microbiome dataset identified families Sphingomonads and *Methylobacteria* as most prevalent, with most reads (> 50%) belonging to *Sphingomonas*. Similarly, a separate study reported these genera as dominant members of the phyllosphere microbiota in two maize genotypes (Kong et al. 2020). Both genera are commonly abundant in the plant phyllosphere, and *Methylobacterium* species can contribute to plant growth (Vacher et al. 2016).

Integrating omics can improve our understanding of plant-microbe interactions, which has been recently reviewed for crop microbiome studies (Kimotho and Maina 2024). Amplicon sequencing and host metabolomics were previously conducted in roots that identified plant coumarins as involved in impacting microbial communities in *Arabidopsis* (Voges et al. 2019). Using metabolomics, transcriptomics, and metataxonomics, another study identified microbial communities associated with different changes in the systemic induction of plant metabolites (Korenblum et al. 2020). A recent study was performed integrating transcriptomics and amplicon sequencing of rhizosphere bacteria in *Asparagus conchinchinensis* (Zhang et al. 2025). The interactions between plants, microbiota, and soil have been extensively studied in the context of the plant rhizosphere. A recent review pinpoints the diurnal rhythmicity of plants, the microbiota, and potential associations between plant physiology and microbial processes and highlights several knowledge gaps that exist in understanding such associations in the rhizosphere (Bending et al. 2025). However, to date, no study has investigated the relationship between plant transcriptome and microbiome across contrasting diurnal periods in leaves. In this study, we addressed this gap by performing pairwise association analyses (cross-correlations) between host transcriptomic profiles and microbial community data (OTU abundances) in maize leaves. Potential symbiotic relationships between microorganisms and the plant have been raised and recently reviewed (Demmig-Adams et al. 2022), which include additional sinks for carbohydrates, supply of nutrients supporting the growth of sink tissues without disrupting nitrogen metabolism, production of regulators of plant metabolism and growth contributing to restoring redox homeostasis.

We identified the expression of several transcription factors cross-correlated with OTU abundances. Interestingly, we also identified two of the circadian cycle genes, GIGANTEA - GI2 and the Pseudo-Response Regulator gene (PRR73 - Zm00001eb397240), having correlated expression with OTUs. The latter family (PRR) is known to be involved in the repression of CCA1/LHY during the day (Lai et al. 2020). This observed association between microbial abundance and expression of some of the circadian cycle genes in leaves is consistent with previous studies in the rhizosphere that reported changes in microbial composition associated with host plants with mutations in circadian cycle genes (Newman et al. 2022; Hubbard et al. 2018).

In our study, we identified an OTU affiliated with *Deinococcus* that was classified as a keystone species and showed significant correlations with the expression of several genes in both day and night networks. *Deinococcus* species are known for their resistance to extreme radiation and oxidative stress (Lim et al. 2019) and have also been reported in the promotion of plant growth (Chitara et al. 2024). Given the harsh conditions of maize leaves, including the day incidence of radiation, this OTU may play an important role in the microbial communities and impact maize health.

## Conclusion

In summary, our integrated midday–midnight analysis of more than 200 maize genotypes establishes that the phyllosphere is orchestrated by a finely tuned temporal interplay between host and microbiome: a pronounced transcriptomic reprogramming, day-time enrichment of photosynthesis and heat-response modules and night-time up-regulation of transcriptional regulators, is mirrored by a subtler but significant phylogenetic restructuring of the bacterial community, whose overall diversity remains stable even as its composition shifts. Keystone taxa occupy distinct temporal niches, with a stress-resistant *Deinococcus* acting as a constant hub, while *Pseudomonas* and *Kineococcus* assume hub roles exclusively by day and night, respectively. Crucially, cross-kingdom correlations link bacterial abundances to core circadian genes such as *GIGANTEA* and *PRR73*, implying that the plant’s internal clock helps shape its leaf microbiome and that key microbes, notably *Deinococcus*, reciprocate irrespective of time of day. These insights provide the first molecular evidence that circadian regulation and microbial ecology are intertwined in the maize phyllosphere, and they lay the groundwork for mechanistic studies, combining host mutants, meta-omics, metabolomics, and synthetic communities, to harness this diurnal duet for improved crop resilience and productivity.

## Acknowledgements

RACS was supported by the Brazilian São Paulo Research Foundation (FAPESP) grant numbers #2021/11057-0 and #2023/11133-3. This work was also supported by FAPESP grants #2014/50884-5 (National Institute of Science and Technology of Bioethanol) and #2020/15230-5 (RCG2I), and CNPq #465319/2014-9. DMRP acknowledges a fellowship from National Council for Scientific and Technological Development – CNPq (311558/2021-6).

## Author contributions

RACS - project administration, funding acquisition, investigation, conceptualization, data curation, formal analysis, software, writing - original draft

KHM - conceptualization, writing - review and editing

JMMP - methodology, data curation, formal analysis, software, writing - review and editing

DJL - visualization, writing - review and editing

CL - methodology, conceptualization, visualization

LWM - conceptualization, supervision, writing - review and editing

DMRP - project administration, funding acquisition, conceptualization, methodology, supervision, resources, writing - review and editing, visualization

JGW - project administration, investigation, data curation, conceptualization, methodology, supervision, resources, writing - review and editing, visualization

### Open Research

#### Code availability

https://github.com/labbces/maize_transcriptome_microbiome_networks

### Supporting information

#### Supplementary Figures

**Sup Fig S1.** Principal Component Analysis (PCA) based on gene expression highlighting maize subpopulations.

**Sup Fig S2.** Co-expression clusters responding to circadian periods. Distribution of gene expression (z-score of log(TPM + 1) values) for co-expression clusters identified as responding to circadian periods.

**Sup Fig S3.** Most frequent orders and genera (GTDB) in the OTU subset that was employed in microbiome analyses and cross-correlations with the maize transcriptome.

**Sup Fig S4.** Tests for differences between diurnal periods in alpha diversity based on (A) evenness, (B) Faith’s PD, and (C) Shannon. P-values (Kruskal-Wallis test) were considered high in all tests (diversity not significantly different).

**Sup Fig S5.** Cross-correlations between transcriptome and bacteriome in maize leaves, determined based on SparXCC permutation values. Distribution of positive (blue) and negative (red) significant cross-correlations identified in day and night samples. Non-significant correlations (near 0) are excluded from the plot.

#### Supplementary Tables

**Sup Table S1.** Expression of maize genes in two diurnal periods and their associated co-expression modules.

**Sup Table S2.** GTDB taxonomy associated with bacterial OTUs.

**Sup Table S3.** Differential abundance tests of bacterial OTUs between midday and midnight diurnal periods.

**Sup Table S4.** Louvain modules and Zi/Pi scores for bacterial OTUs in two diurnal periods.

**Sup Table S5.** Cross-correlations between bacterial OTUs and maize gene expression in two diurnal periods.

**Sup Table S6.** Annotation of Transcription Factor Families (TFF), Other Transcriptional Regulators (OTR), and Orphans in cross-correlated maize genes in two diurnal periods.

## Notes

https://github.com/SantosRAC/maize_transcriptome_microbiome_diurnal_periods/

### Competing Interest Statement

The authors have declared no competing interest.

## References

Abadi, Vahid Alah Jahandideh Mahjen, Mozhgan Sepehri, Hadi Asadi Rahmani, Mehdi Zarei, Abdolmajid Ronaghi, Seyed Mohsen Taghavi, and Mahdieh Shamshiripour. 2020. “Role of Dominant Phyllosphere Bacteria with Plant Growth–promoting Characteristics on Growth and Nutrition of Maize (Zea Mays L.).” Journal of Soil Science and Plant Nutrition 20 (4): 2348–63.

Banerjee, Samiran, Klaus Schlaeppi, and Marcel G. A. van der Heijden. 2018. “Keystone Taxa as Drivers of Microbiome Structure and Functioning.” Nature Reviews. Microbiology 16 (9): 567–76.

Becker, Martin, Huda Nassar, Camilo Espinosa, Ina A. Stelzer, Dorien Feyaerts, Eloise Berson, Neda H. Bidoki, et al. 2023. “Large-Scale Correlation Network Construction for Unraveling the Coordination of Complex Biological Systems.” Nature Computational Science 3 (4): 346–59.

Bending, Gary D., Amy Newman, Emma Picot, Ryan M. Mushinski, Davey L. Jones, and Isabelle A. Carré. 2025. “Diurnal Rhythmicity in the Rhizosphere Microbiome-Mechanistic Insights and Significance for Rhizosphere Function.” Plant, Cell & Environment 48 (3): 2040–52.

Bendix, Claire, Carine M. Marshall, and Frank G. Harmon. 2015. “Circadian Clock Genes Universally Control Key Agricultural Traits.” Molecular Plant 8 (8): 1135–52.

Bolyen, Evan, Jai Ram Rideout, Matthew R. Dillon, Nicholas A. Bokulich, Christian C. Abnet, Gabriel A. Al-Ghalith, Harriet Alexander, et al. 2019. “Reproducible, Interactive, Scalable and Extensible Microbiome Data Science Using QIIME 2.” Nature Biotechnology 37 (8): 852–57.

Borba, Ana Rita, Ivan Reyna-Llorens, Patrick J. Dickinson, Gareth Steed, Paulo Gouveia, Alicja M. Górska, Celia Gomes, et al. 2023. “Compartmentation of Photosynthesis Gene Expression in C4 Maize Depends on Time of Day.” Plant Physiology 193 (4): 2306–20.

Chen, Z. Jeffrey. 2013. “Genomic and Epigenetic Insights into the Molecular Bases of Heterosis.” Nature Reviews. Genetics 14 (7): 471–82.

Chitara, Manoj Kumar, Rajesh Pratap Singh, Narendra Kumar Singh, Yogendra Singh Rajpurohit, and Hari S. Misra. 2024. “Plant Growth-Promoting Potential of Deinococci Spp. Evaluated Using Zea Mays and Lens Culinaris Crops.” Journal of Plant Growth Regulation 43 (11): 4384–95.

Csárdi, Gábor, Tamás Nepusz, Kirill Müller, Szabolcs Horvát, Vincent Traag, Fabio Zanini, and Daniel Noom. 2025. Igraph for R: R Interface of the Igraph Library for Graph Theory and Network Analysis. Zenodo. 10.5281/ZENODO.7682609.

Demmig-Adams, B., S. K. Polutchko, M. C. Zenir, P. Fourounjian, J. J. Stewart, M. López-Pozo, and W. W. Adams 3rd. 2022. “Intersections: Photosynthesis, Abiotic Stress, and the Plant Microbiome.” Photosynthetica 60 (1): 59–69.

Dodd, Antony N., Neeraj Salathia, Anthony Hall, Eva Kévei, Réka Tóth, Ferenc Nagy, Julian M. Hibberd, Andrew J. Millar, and Alex A. R. Webb. 2005. “Plant Circadian Clocks Increase Photosynthesis, Growth, Survival, and Competitive Advantage.” Science (New York, N.Y.) 309 (5734): 630–33.

Douglas, Gavin M., Vincent J. Maffei, Jesse R. Zaneveld, Svetlana N. Yurgel, James R. Brown, Christopher M. Taylor, Curtis Huttenhower, and Morgan G. I. Langille. 2020. “PICRUSt2 for Prediction of Metagenome Functions.” Nature Biotechnology 38 (6): 685–88.

Edgar, Rachel S., Edward W. Green, Yuwei Zhao, Gerben van Ooijen, Maria Olmedo, Ximing Qin, Yao Xu, et al. 2012. “Peroxiredoxins Are Conserved Markers of Circadian Rhythms.” Nature 485 (7399): 459–64.

Gao, Zhi-Fang, Xiu Yang, Yingchang Mei, Jiao Zhang, Qing Chao, and Bai-Chen Wang. 2023. “A Dynamic Phosphoproteomic Analysis Provides Insight into the C4 Plant Maize (Zea Mays L.) Response to Natural Diurnal Changes.” The Plant Journal : For Cell and Molecular Biology 113 (2): 291–307.

Greenham, Kathleen, and C. Robertson McClung. 2015. “Integrating Circadian Dynamics with Physiological Processes in Plants.” Nature Reviews. Genetics 16 (10): 598–610.

Guimerà, Roger, and Luís A. Nunes Amaral. 2005. “Functional Cartography of Complex Metabolic Networks.” Nature 433 (7028): 895–900.

Gunnigle, Eoin, Aline Frossard, Jean-Baptiste Ramond, Leandro Guerrero, Mary Seely, and Don A. Cowan. 2017. “Diel-Scale Temporal Dynamics Recorded for Bacterial Groups in Namib Desert Soil.” Scientific Reports 7 (January):40189.

Harmer, Stacey L. 2009. “The Circadian System in Higher Plants.” Annual Review of Plant Biology 60:357–77.

Haydon, Michael J., Ángela Román, and Waheed Arshad. 2015. “Nutrient Homeostasis within the Plant Circadian Network.” Frontiers in Plant Science 6 (April):299.

Hayes, Kevin R., Mary Beatty, Xin Meng, Carl R. Simmons, Jeffrey E. Habben, and Olga N. Danilevskaya. 2010. “Maize Global Transcriptomics Reveals Pervasive Leaf Diurnal Rhythms but Rhythms in Developing Ears Are Largely Limited to the Core Oscillator.” PloS One 5 (9): e12887.

Hidalgo-Martinez, Kelly, Admir José Giachini, Marcio Schneider, Adriana Soriano, Marcus Paulus Baessa, Luiz Fernando Martins, and Valéria Maia de Oliveira. 2024. “Shifts in Structure and Dynamics of the Soil Microbiome in Biofuel/fuel Blend-Affected Areas Triggered by Different Bioremediation Treatments.” Environmental Science and Pollution Research International 31 (23): 33663–84.

Hubbard, Charley J., Marcus T. Brock, Linda Ta van Diepen, Loïs Maignien, Brent E. Ewers, and Cynthia Weinig. 2018. “The Plant Circadian Clock Influences Rhizosphere Community Structure and Function.” The ISME Journal 12 (2): 400–410.

Hufford, Matthew B., Arun S. Seetharam, Margaret R. Woodhouse, Kapeel M. Chougule, Shujun Ou, Jianing Liu, William A. Ricci, et al. 2021. “De Novo Assembly, Annotation, and Comparative Analysis of 26 Diverse Maize Genomes.” Science (New York, N.Y.) 373 (6555): 655–62.

Jensen, Ib Thorsgaard, Luc Janss, Simona Radutoiu, and Rasmus Waagepetersen. 2024. “Compositionally Aware Estimation of Cross-Correlations for Microbiome Data.” PloS One 19 (6): e0305032.

Jones, Philip, David Binns, Hsin-Yu Chang, Matthew Fraser, Weizhong Li, Craig McAnulla, Hamish McWilliam, et al. 2014. “InterProScan 5: Genome-Scale Protein Function Classification.” Bioinformatics (Oxford, England) 30 (9): 1236–40.

Katoh, Kazutaka, and Hiroyuki Toh. 2008. “Recent Developments in the MAFFT Multiple Sequence Alignment Program.” Briefings in Bioinformatics 9 (4): 286–98.

Kimotho, Roy Njoroge, and Solomon Maina. 2024. “Unraveling Plant-Microbe Interactions: Can Integrated Omics Approaches Offer Concrete Answers?” Journal of Experimental Botany 75 (5): 1289–1313.

Klopfenstein, D. V., Liangsheng Zhang, Brent S. Pedersen, Fidel Ramírez, Alex Warwick Vesztrocy, Aurélien Naldi, Christopher J. Mungall, et al. 2018. “GOATOOLS: A Python Library for Gene Ontology Analyses.” Scientific Reports 8 (1): 10872.

Ko, Dae Kwan, Dominica Rohozinski, Qingxin Song, Samuel H. Taylor, Thomas E. Juenger, Frank G. Harmon, and Z. Jeffrey Chen. 2016. “Temporal Shift of Circadian-Mediated Gene Expression and Carbon Fixation Contributes to Biomass Heterosis in Maize Hybrids.” PLoS Genetics 12 (7): e1006197.

Kolde, Raivo. 2025 “pheatmap: Pretty Heatmaps” https://github.com/raivokolde/pheatmap

Kong, Xiao, Zhenfei Han, Xin Tai, Decai Jin, Sen Ai, Xiaoxu Zheng, and Zhihui Bai. 2020. “Maize (Zea Mays L. Sp.) Varieties Significantly Influence Bacterial and Fungal Community in Bulk Soil, Rhizosphere Soil and Phyllosphere.” FEMS Microbiology Ecology 96 (3). 10.1093/femsec/fiaa020.

Korenblum, Elisa, Yonghui Dong, Jedrzej Szymanski, Sayantan Panda, Adam Jozwiak, Hassan Massalha, Sagit Meir, Ilana Rogachev, and Asaph Aharoni. 2020. “Rhizosphere Microbiome Mediates Systemic Root Metabolite Exudation by Root-to-Root Signaling.” Proceedings of the National Academy of Sciences of the United States of America 117 (7): 3874–83.

Kremling, Karl A. G., Shu-Yun Chen, Mei-Hsiu Su, Nicholas K. Lepak, M. Cinta Romay, Kelly L. Swarts, Fei Lu, Anne Lorant, Peter J. Bradbury, and Edward S. Buckler. 2018. “Dysregulation of Expression Correlates with Rare-Allele Burden and Fitness Loss in Maize.” Nature 555 (7697): 520–23.

Krueger, Felix, Frankie James, Phil Ewels, Ebrahim Afyounian, Michael Weinstein, Benjamin Schuster-Boeckler, Gert Hulselmans, and sclamons. 2023. FelixKrueger/TrimGalore: v0.6.10 - Add Default Decompression Path. Zenodo. 10.5281/ZENODO.7598955.

Kurtz, Zachary D., Christian L. Müller, Emily R. Miraldi, Dan R. Littman, Martin J. Blaser, and Richard A. Bonneau. 2015. “Sparse and Compositionally Robust Inference of Microbial Ecological Networks.” PLoS Computational Biology 11 (5): e1004226.

Lai, Xianjun, Claire Bendix, Lang Yan, Yang Zhang, James C. Schnable, and Frank G. Harmon. 2020. “Interspecific Analysis of Diurnal Gene Regulation in Panicoid Grasses Identifies Known and Novel Regulatory Motifs.” BMC Genomics 21 (1): 428.

Li, Chenxin, Natalie C. Deans, and C. Robin Buell. 2023. “‘Simple Tidy GeneCoEx’: A Gene Co-Expression Analysis Workflow Powered by Tidyverse and Graph-Based Clustering in R.” The Plant Genome 16 (2): e20323.

Lim, Sangyong, Jong-Hyun Jung, Laurence Blanchard, and Arjan de Groot. 2019. “Conservation and Diversity of Radiation and Oxidative Stress Resistance Mechanisms in Deinococcus Species.” FEMS Microbiology Reviews 43 (1): 19–52.

Lin, Huang, and Shyamal Das Peddada. 2020. “Analysis of Compositions of Microbiomes with Bias Correction.” Nature Communications 11 (1): 3514.

Lin, Tiantian, Jiayao Tang, Shuying Li, Shujiang Li, Shan Han, Yinggao Liu, Chunlin Yang, Gang Chen, Lianghua Chen, and Tianhui Zhu. 2023. “Drought Stress-Mediated Differences in Phyllosphere Microbiome and Associated Pathogen Resistance between Male and Female Poplars.” The Plant Journal : For Cell and Molecular Biology 115 (4): 1100–1113.

Li, Zhi, Andan Zhu, Qingxin Song, Helen Y. Chen, Frank G. Harmon, and Z. Jeffrey Chen. 2020. “Temporal Regulation of the Metabolome and Proteome in Photosynthetic and Photorespiratory Pathways Contributes to Maize Heterosis.” The Plant Cell 32 (12): 3706–22.

Matchado, Monica Steffi, Michael Lauber, Sandra Reitmeier, Tim Kacprowski, Jan Baumbach, Dirk Haller, and Markus List. 2021. “Network Analysis Methods for Studying Microbial Communities: A Mini Review.” Computational and Structural Biotechnology Journal 19 (May):2687–98.

“Microbial Interaction-Driven Community Differences as Revealed by Network Analysis.” 2021. Computational and Structural Biotechnology Journal 19 (January):6000–6008.

Minh, Bui Quang, Heiko A. Schmidt, Olga Chernomor, Dominik Schrempf, Michael D. Woodhams, Arndt von Haeseler, and Robert Lanfear. 2020. “IQ-TREE 2: New Models and Efficient Methods for Phylogenetic Inference in the Genomic Era.” Molecular Biology and Evolution 37 (5): 1530–34.

Mistry, Jaina, Robert D. Finn, Sean R. Eddy, Alex Bateman, and Marco Punta. 2013. “Challenges in Homology Search: HMMER3 and Convergent Evolution of Coiled-Coil Regions.” Nucleic Acids Research 41 (12): e121.

Newman, Amy, Emma Picot, Sian Davies, Sally Hilton, Isabelle A. Carré, and Gary D. Bending. 2022. “Circadian Rhythms in the Plant Host Influence Rhythmicity of Rhizosphere Microbiota.” BMC Biology 20 (1): 235.

Ni, Zhongfu, Eun-Deok Kim, Misook Ha, Erika Lackey, Jianxin Liu, Yirong Zhang, Qixin Sun, and Z. Jeffrey Chen. 2009. “Altered Circadian Rhythms Regulate Growth Vigour in Hybrids and Allopolyploids.” Nature 457 (7227): 327–31.

Olesen, Jens M., Jordi Bascompte, Yoko L. Dupont, and Pedro Jordano. 2007. “The Modularity of Pollination Networks.” Proceedings of the National Academy of Sciences of the United States of America 104 (50): 19891–96.

Palberg, Daniel, Anna Kisiata, Gabriel Lemes Jorge, and R. J. Neil Emery. 2022. “A survey of Methylobacterium species and strains reveals widespread production and varying profiles of cytokinin phytohormones”. BMC Microbiology 22 (1): 49.

Parks, Donovan H., Maria Chuvochina, Christian Rinke, Aaron J. Mussig, Pierre-Alain Chaumeil, and Philip Hugenholtz. 2022. “GTDB: An Ongoing Census of Bacterial and Archaeal Diversity through a Phylogenetically Consistent, Rank Normalized and Complete Genome-Based Taxonomy.” Nucleic Acids Research 50 (D1): D785–94.

Patro, Rob, Geet Duggal, Michael I. Love, Rafael A. Irizarry, and Carl Kingsford. 2017. “Salmon Provides Fast and Bias-Aware Quantification of Transcript Expression.” Nature Methods 14 (4): 417–19.

Pérez-Rodríguez, Paulino, Diego Mauricio Riaño-Pachón, Luiz Gustavo Guedes Corrêa, Stefan A. Rensing, Birgit Kersten, and Bernd Mueller-Roeber. 2010. “PlnTFDB: Updated Content and New Features of the Plant Transcription Factor Database.” Nucleic Acids Research 38 (Database issue): D822–27.

Robeson, Michael S., 2nd, Devon R. O’Rourke, Benjamin D. Kaehler, Michal Ziemski, Matthew R. Dillon, Jeffrey T. Foster, and Nicholas A. Bokulich. 2021. “RESCRIPt: Reproducible Sequence Taxonomy Reference Database Management.” PLoS Computational Biology 17 (11): e1009581.

Sanchez, Sabrina E., and Steve A. Kay. 2016. “The Plant Circadian Clock: From a Simple Timekeeper to a Complex Developmental Manager.” Cold Spring Harbor Perspectives in Biology 8 (12). 10.1101/cshperspect.a027748.

Santos-Medellín, Christian, Joseph Edwards, Bao Nguyen, and Venkatesan Sundaresan. 2022. “Acquisition of a Complex Root Microbiome Reshapes the Transcriptomes of Rice Plants.” The New Phytologist 235 (5): 2008–21.

Sohrabi, Reza, Bradley C. Paasch, Julian A. Liber, and Sheng Yang He. 2023. “Phyllosphere Microbiome.” Annual Review of Plant Biology 74 (May):539–68.

Staley, Christopher, Abigail P. Ferrieri, Malak M. Tfaily, Yaya Cui, Rosalie K. Chu, Ping Wang, Jared B. Shaw, et al. 2017. “Diurnal Cycling of Rhizosphere Bacterial Communities Is Associated with Shifts in Carbon Metabolism.” Microbiome 5 (1): 65.

Vacher, Corinne, Arndt Hampe, Annabel J. Porté, Ursula Sauer, Stéphane Compant, and Cindy E. Morris. 2016. “The Phyllosphere: Microbial Jungle at the Plant–climate Interface.” Annual Review of Ecology, Evolution, and Systematics 47 (1): 1–24.

Varoquaux, Nelle, Benjamin Cole, Cheng Gao, Grady Pierroz, Christopher R. Baker, Dhruv Patel, Mary Madera, et al. 2019. “Transcriptomic Analysis of Field-Droughted Sorghum from Seedling to Maturity Reveals Biotic and Metabolic Responses.” Proceedings of the National Academy of Sciences of the United States of America 116 (52): 27124–32.

Voges, Mathias J. E. E. E., Yang Bai, Paul Schulze-Lefert, and Elizabeth S. Sattely. 2019. “Plant-Derived Coumarins Shape the Composition of an Synthetic Root Microbiome.” Proceedings of the National Academy of Sciences of the United States of America 116 (25): 12558–65.

Wallace, Jason G., Karl A. Kremling, Lynsey L. Kovar, and Edward S. Buckler. 2018. “Quantitative Genetics of the Maize Leaf Microbiome.” Phytobiomes Journal 2 (4): 208–24.

Watson, Christopher G. 2015. “BrainGraph: Graph Theory Analysis of Brain MRI Data.” CRAN: Contributed Packages. The R Foundation. 10.32614/cran.package.braingraph.

Xiong, Xiang, Yanfang Xing, Jinzhi He, Li Wang, Zhenzhen Shen, Wenli Chen, and Qiaoyun Huang. 2021. “Keystone Species Determine the ‘Selection Mechanism’ of Multispecies Biofilms for Bacteria from Soil Aggregates.” The Science of the Total Environment 773 (June):145069.

Xu, Ling, Zhaobin Dong, Dawn Chiniquy, Grady Pierroz, Siwen Deng, Cheng Gao, Spencer Diamond, et al. 2021. “Genome-Resolved Metagenomics Reveals Role of Iron Metabolism in Drought-Induced Rhizosphere Microbiome Dynamics.” Nature Communications 12 (1): 3209.

Xu, Ling, Dan Naylor, Zhaobin Dong, Tuesday Simmons, Grady Pierroz, Kim K. Hixson, Young-Mo Kim, et al. 2018. “Drought Delays Development of the Sorghum Root Microbiome and Enriches for Monoderm Bacteria.” Proceedings of the National Academy of Sciences of the United States of America 115 (18): E4284–93.

Xu, Nuohan, Qianqiu Zhao, Zhenyan Zhang, Qi Zhang, Yan Wang, Guoyan Qin, Mingjing Ke, et al. 2022. “Phyllosphere Microorganisms: Sources, Drivers, and Their Interactions with Plant Hosts.” Journal of Agricultural and Food Chemistry 70 (16): 4860–70.

Xu, Xiaodong, Li Yuan, Xin Yang, Xiao Zhang, Lei Wang, and Qiguang Xie. 2022. “Circadian Clock in Plants: Linking Timing to Fitness.” Journal of Integrative Plant Biology 64 (4): 792–811.

Yang, Yaodong, Mumtaz Ali Saand, Liyun Huang, Walid Badawy Abdelaal, Jun Zhang, Yi Wu, Jing Li, Muzafar Hussain Sirohi, and Fuyou Wang. 2021. “Applications of Multi-Omics Technologies for Crop Improvement.” Frontiers in Plant Science 12 (September):563953.

Yu, Guangchuang, Li-Gen Wang, Yanyan Han, and Qing-Yu He. 2012. “clusterProfiler: An R Package for Comparing Biological Themes among Gene Clusters.” Omics : A Journal of Integrative Biology 16 (5): 284–87.

Zhan, Chengfang, Haruna Matsumoto, Yufei Liu, and Mengcen Wang. 2022. “Pathways to Engineering the Phyllosphere Microbiome for Sustainable Crop Production.” Nature Food 3 (12): 997–1004.

Zhang, Xiaoyong, Shuai Yang, Jingsheng Yu, Xiongwei Liu, Xuebo Tang, Liuyan Wang, Jinglan Chen, et al. 2025. “Integrated Amplicon Sequencing and Transcriptomic Sequencing Technology Reveals Changes in the Bacterial Community and Gene Expression in the Rhizosphere Soil of *Asparagus Cochinchinensis*.” Medicinal Plant Biology 4 (1): 0–0.

Zhang, Yuanyuan, Yan Wang, Hua Wei, Na Li, Wenwen Tian, Kang Chong, and Lei Wang. 2018. “Circadian Evening Complex Represses Jasmonate-Induced Leaf Senescence in Arabidopsis.” Molecular Plant 11 (2): 326–37.

Zhao, Kankan, Bin Ma, Yan Xu, Erinne Stirling, and Jianming Xu. 2021. “Light Exposure Mediates Circadian Rhythms of Rhizosphere Microbial Communities.” The ISME Journal 15 (9): 2655–64.

Zhao, Yongping, Binbin Zhao, Yurong Xie, Hong Jia, Yongxiang Li, Miaoyun Xu, Guangxia Wu, et al. 2023. “The Evening Complex Promotes Maize Flowering and Adaptation to Temperate Regions.” The Plant Cell 35 (1): 369–89.

Zhu, Guofan, Ruijun Du, Daolin Du, Jiazhong Qian, and Mao Ye. 2021. “Keystone Taxa Shared between Earthworm Gut and Soil Indigenous Microbial Communities Collaboratively Resist Chlordane Stress.” Environmental Pollution (Barking, Essex : 1987) 283 (August):117095.

Zhu, Yong-Guan, Jingjing Peng, Cai Chen, Chao Xiong, Shule Li, Anhui Ge, Ertao Wang, and Werner Liesack. 2023. “Harnessing Biological Nitrogen Fixation in Plant Leaves.” Trends in Plant Science 28 (12): 1391–1405.

