## Supplementary Figures for "Identifying associations between maize leaf transcriptome and bacteriome during different diurnal periods"

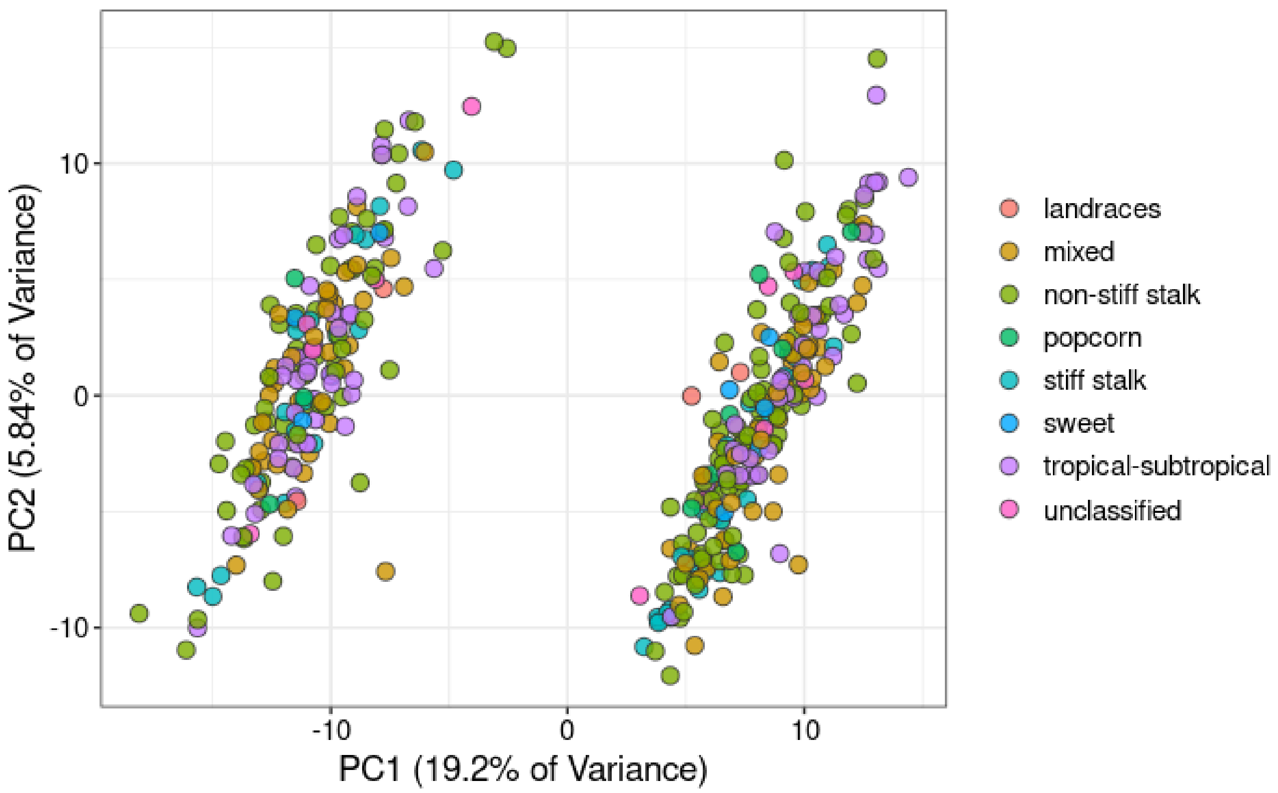


**Sup Fig S1.** Principal Component Analysis (PCA) based on gene expression highlighting maize subpopulations.

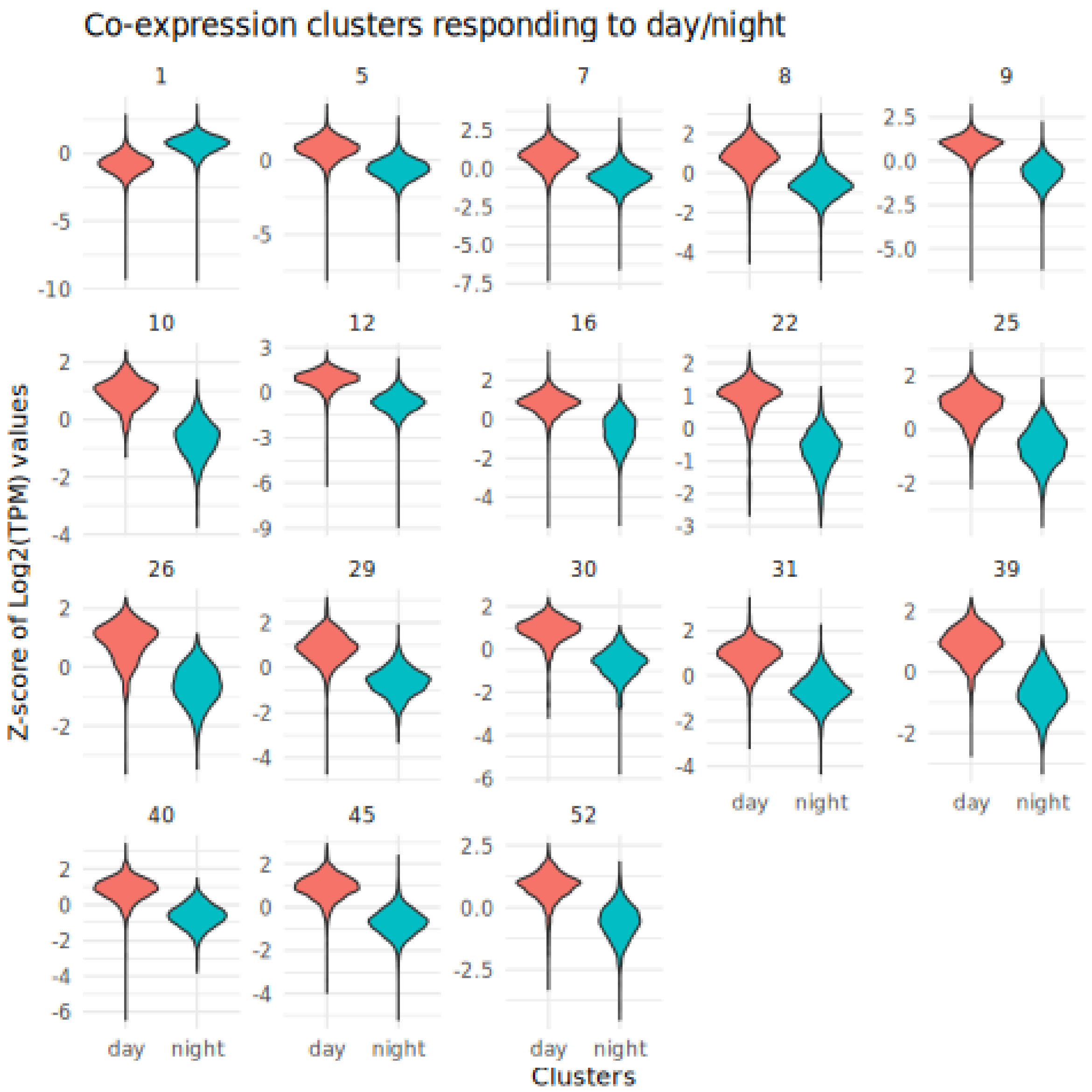


**Sup Fig S2.** Co-expression clusters responding to circadian periods. Distribution of gene expression (z-score of log(TPM + 1) values) for co-expression clusters identified as responding to circadian periods.

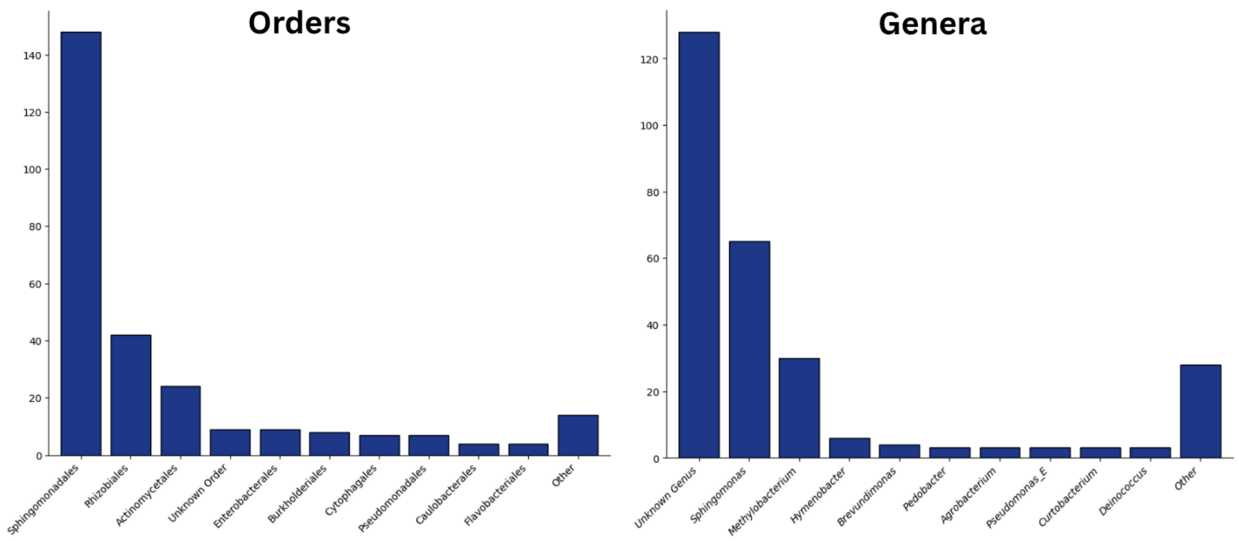


**Sup Fig S3.** Most frequent orders and genera (GTDB) in the OTU subset that was employed in microbiome analyses and cross-correlations with the maize transcriptome.

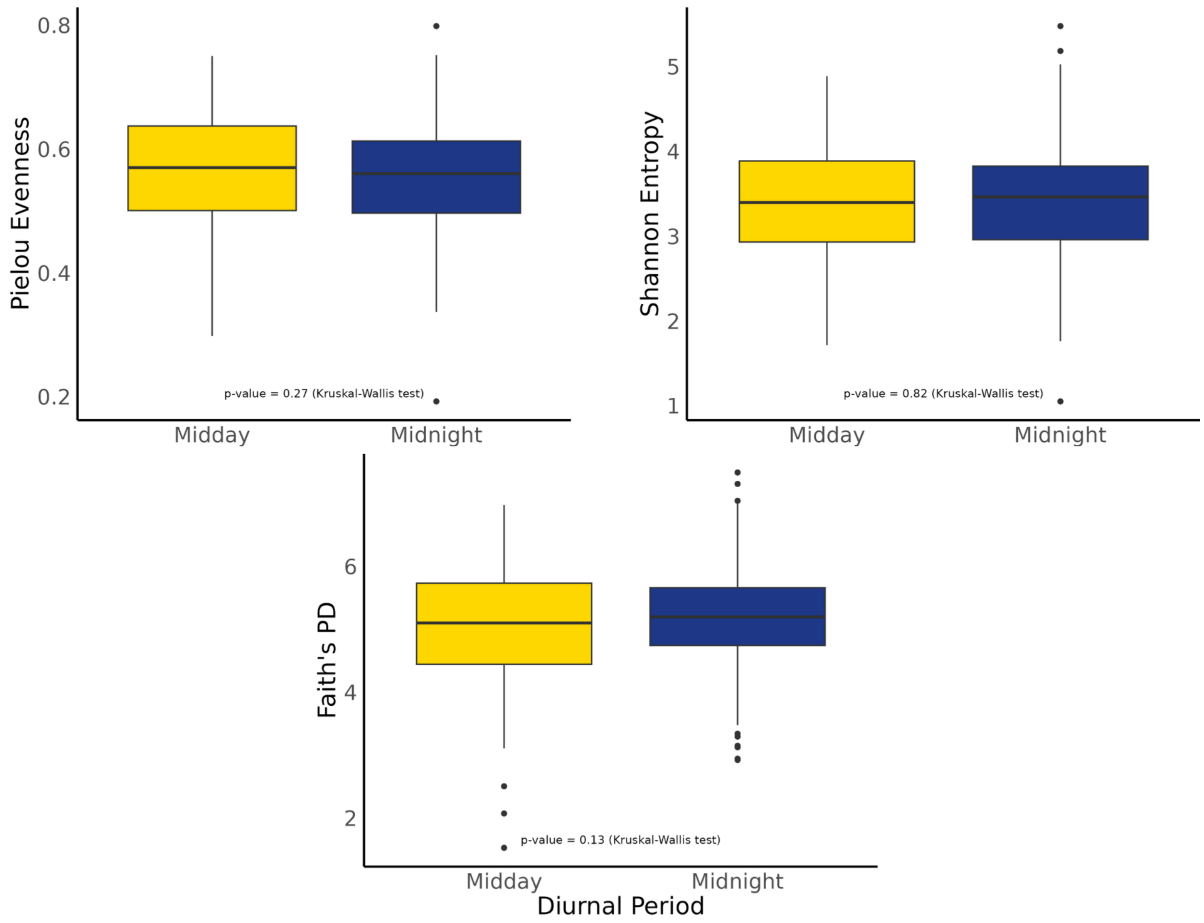


**Sup Fig S4.** Tests for differences between diurnal periods in alpha diversity based on (A) evenness, (B) Faith's PD, and (C) Shannon. P-values (Kruskal-Wallis test) were considered high in all tests (diversity not significantly different).

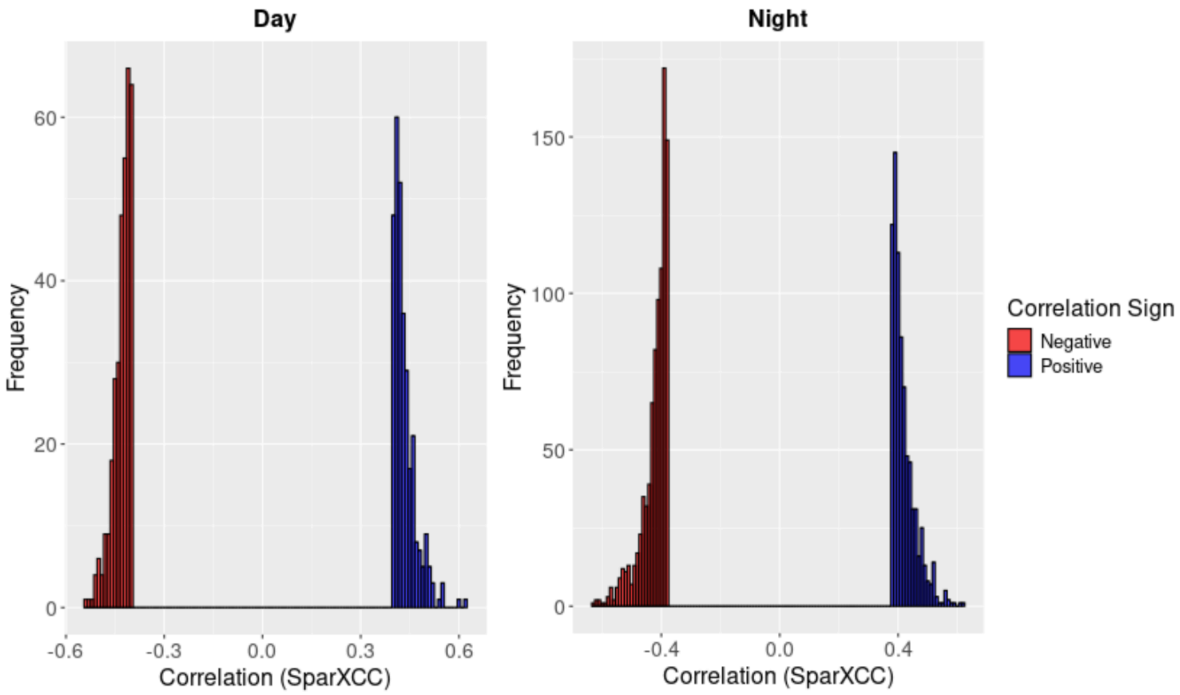


**Sup Fig S5.** Cross-correlations between transcriptome and bacteriome in maize leaves, determined based on SparXCC permutation values. Distribution of positive (blue) and negative (red) significant cross-correlations identified in day and night samples. Non-significant correlations (near 0) are excluded from the plot.
